# Megalin deficiency perturbs retinal homeostasis and impairs cathepsin D processing and phagosome-lysosome maturation in the retinal pigment epithelium

**DOI:** 10.64898/2026.06.26.734689

**Authors:** Ditte K. Rasmussen, Pernille L. Marschall, Soo Hyeon Lee, Tina Storm, Thomas S. Jakobsen, Qi Wu, Anne Louise Askou, Robert A. Fenton, Thomas Corydon, Vinit B. Mahajan, Rikke Nielsen

**Author notes:** Co-first authors (Ditte K. Rasmussen, Pernille L. Marschall). Co-corresponding authors: **Lead contact:** Rikke Nielsen, Ph.D., Department of Biomedicine, Aarhus University, Aarhus, Denmark. Requests for further information and resources should be directed to and will be fulfilled by the lead contact, Vinit B. Mahajan M.D., Ph.D., Byers Eye Institute, Department of Ophthalmology, Stanford University, Palo Alto, CA, 94304 USA.

## Abstract

The multiligand endocytic receptor, megalin (LRP2), is expressed in the retinal pigment epithelium (RPE) and patients lacking the receptor develop high myopia. Despite its established role in retinal development, the contribution of megalin to retinal homeostasis in the normally developed/mature eye remains poorly understood. Here, we investigated megalin function using an inducible knockout mouse (KO) model and human iPSC-derived RPE with megalin knockdown (KD) to distinguish post-developmental homeostatic functions from developmental effects. *In vivo, m*egalin ablation caused progressive retinal degeneration and visual impairment, with morphological abnormalities in the RPE but no changes in myopia-associated ocular phenotypes including axial length and intraocular pressure. Proteomic profiling of megalin-KO RPE revealed reduction of autophagy-related proteins. In line with this, megalin deficiency was associated with accumulation of pro-cathepsin D, and perturbed rhodopsin turnover. This was supported *in vitro*, where trafficking of photoreceptor outer segment (POS) containing phagosomes to lysosomes was reduced, suggesting disturbed phagosome maturation. Megalin KD did not measurably impair initial uptake of POS discs, but delayed rhodopsin degradation, indicating defective post-ingestion processing. Together, these findings establish megalin as a key regulator of retinal homeostasis in the mature eye by controlling phagosome-lysosome fusion in the RPE and suggest that megalin dysfunction contributes to slowly progressive retinal degeneration. This positions megalin as a potential therapeutic target in lysosomal degenerative diseases in the retina.

## Introduction

The retinal pigment epithelium (RPE) is essential for maintaining retinal structure and function. This polarized epithelial monolayer, located between the photoreceptors and the choroid, constitutes the outer blood-retina barrier and provides metabolic support, regulates nutrient and ion transport, participates in the visual cycle, and facilitates the continuous turnover of photoreceptor outer segment (POS) discs[1]. Each RPE cell interfaces with approximately 50 photoreceptors and phagocytoses roughly 10% of their outer segments daily[2, 3]. This places an exceptional degradative burden on the RPE, making it one of the most phagocytically active cell types in the body[4].

POS disc clearance is a highly coordinated process that integrates phagocytosis with intracellular trafficking and lysosomal degradation. Through MER Proto-Oncogene, Tyrosine Kinase (MerTK) and integrin-mediated signaling, RPE cells form a phagocytic cup that engulfs and internalizes the tip of a POS[5, 6]. Following internalization, phagosomes undergo maturation through recruitment of autophagy- and endo-lysosome-associated machinery, including LC3-associated phagocytosis (LAP) which may facilitate fusion with lysosomes [7–9]. Efficient degradation depends on timely delivery of cargo to mature lysosomes[10]. During lysosomal maturation, progressive acidification promotes cleavage of pro-cathepsin D to its active form, enabling fusion with POS-containing phagosomes and subsequent breakdown and recycling of POS-derived material, such as retinoids[8]. In young, healthy RPE, this process proceeds efficiently; however, in aging or disease, incomplete degradation leads to accumulation of undigested material, reflecting dysfunction of the degradative system[11, 12]. The high demand placed on these pathways renders the RPE particularly sensitive to disruptions in intracellular trafficking and degradation. Consequently, dysfunction of the RPE is a central driver in blinding diseases, such as age-related macular degeneration (AMD) and high myopia, underscoring the importance of identifying molecular regulators of these processes[13, 14].

Megalin (also known as LRP2) is a large multiligand endocytic receptor extensively studied in the kidney, where it cooperates with the cubilin-amnionless complex to mediate reabsorption of numerous proteins from the glomerular filtrate, including cathepsin B[15]. In the eye, megalin expression has been detected in several retinal cell types, but converging evidence indicates that the RPE is its principal site of retinal expression[14, 16, 17]. Within the RPE, megalin localizes to the apical membrane as well as to intracellular vesicular compartments[16, 18].

Complete absence of megalin in humans causes Donnai-Barrow/Facio-oculo-acoustico-renal (DB/FOAR) syndrome, which includes high myopia among its clinical features [16, 19, 20]. Fetal loss of megalin in mice leads to enlarged ocular globes and high myopia mimicking the ocular phenotype in patients [16, 18]. LRP2 variations have also been linked to retinal dystrophy[21]. Recent studies identify megalin as a potential driver of non-syndromic high myopia, further underscoring its role in retinal homeostasis[14]. Mechanistically, this has been linked to downregulation of retinol metabolism and pathways related to solute transmembrane transport in iPSC-derived RPE cells following megalin silencing[14]. Fetal knockout (KO) of megalin is associated with decreased expression of endo-lysosomal proteins, while early postnatal KO of megalin disturbed melanosome formation[16, 18, 22]. Together, these findings suggest that megalin functions not only in apical endocytic uptake from the subretinal space but also in intracellular trafficking and lysosome-related organelle maturation.

Thus, evidence clearly demonstrates that fetal KO of megalin leads to altered ocular development and changes in the endo-lysosomal system. To understand the role of megalin in the adult retina, we sought to determine the alterations of POS disc handling in the RPE using inducible megalin KO animal models and iPSC-derived RPE cells with megalin knockdown (KD).

## Results

### Inducible models enable the study of megalin in the normally developed retina

This study aimed to investigate the role of megalin in the normally developed retina (referred to as mature) rather than the consequences of megalin deficiency during ocular development. To this end, we established both an inducible megalin KO mouse model and a human iPSC-derived RPE model with megalin KD.

In mice, western blotting confirmed efficient ablation of megalin 4 weeks after KO induction (ages in the following refer to time after induction), and RPE flatmount staining demonstrated a mosaic KO pattern with very few megalin-positive cells scattered throughout the RPE layer (Fig. 1A). In parallel, iPSCs from the Personal Genome Project were differentiated into mature RPE monolayers. Differentiation was confirmed by loss of pluripotency markers (OCT4, SOX2) and expression of RPE markers (OTX2, RPE65, CRALBP), along with megalin expression and formation of a polarized hexagonal monolayer marked by ZO-1 and occludin (Fig. 1B). Megalin KD was achieved by screening short hairpin RNAs (shRNAs) and selecting the most efficient construct (shRNA D), which reduced protein levels by approximately 60%, as confirmed by western blotting and immunofluorescence (Fig. 1C).

**Figure 1:**
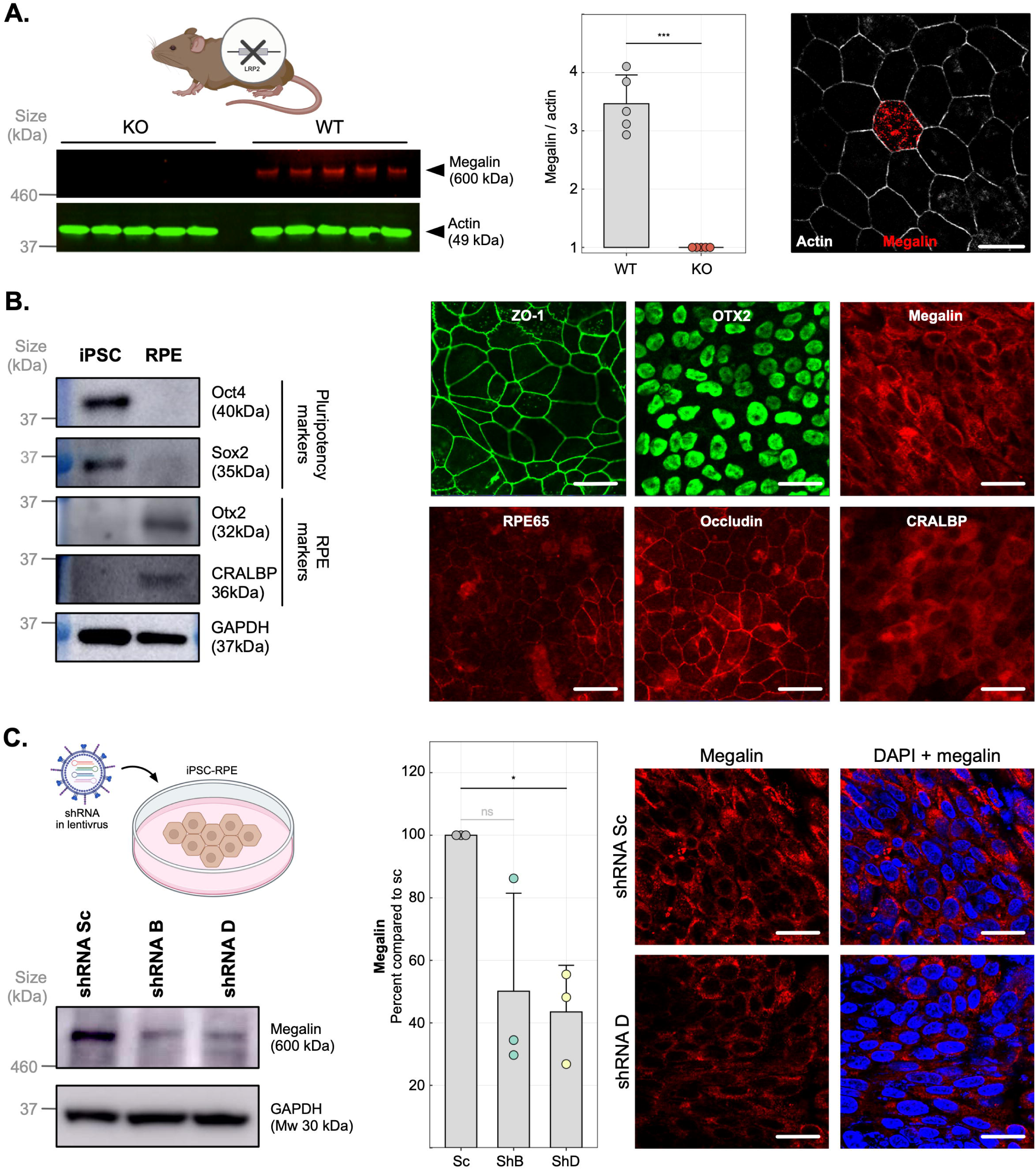
Megalin ablation in mice and iPSC-RPE cells. **A)** Quantitative western blotting for megalin in isolated RPE cell from the inducible mouse model and immunofluorescence of RPE flatmount from the mouse model stained for megalin (red) and actin (white). Scale bar: 20 µm. **B)** Representative western blotting of differentiated iPSC-RPE cells for pluripotency markers OCT4 and SOX2, RPE differentiation markers OTX2, and CRALBP and immunofluorescence for megalin and RPE markers OTX2, RPE65, and CRALBP as well as tight junction markers ZO-1 and Occludin. Scale bars: 50 µm. **C)** Quantitative western blotting for megalin abundance after treatment with short hairpin RNA targeting megalin in differentiated iPSC-RPE cells and representative immunofluorescence of monolayers for megalin (red) and DAPI (blue) in the shRNA treated cells. Scale bars: 50 µm.

Given the difference in KO efficiency between the two systems, we hereafter refer to the mouse model as megalin-KO and the cell model as megalin KD. Together, these models enabled investigation of megalin function in mature RPE physiology.

### Megalin ablation leads to progressive retinal degeneration

To determine how loss of megalin affects retinal integrity over time, we performed longitudinal structural and functional analyses of the megalin KO-mice. Optical coherence tomography (OCT) showed gradual thinning of the retina over 18 months, with the most pronounced changes observed in the outer nuclear layer (ONL) and the inner nuclear layer (INL) (Fig. 2A). Subtle thinning of the outer retina was detectable already within weeks after induction, whereas inner retinal layers were comparatively preserved at early time points and only displayed milder changes later in disease progression.

**Figure 2:**
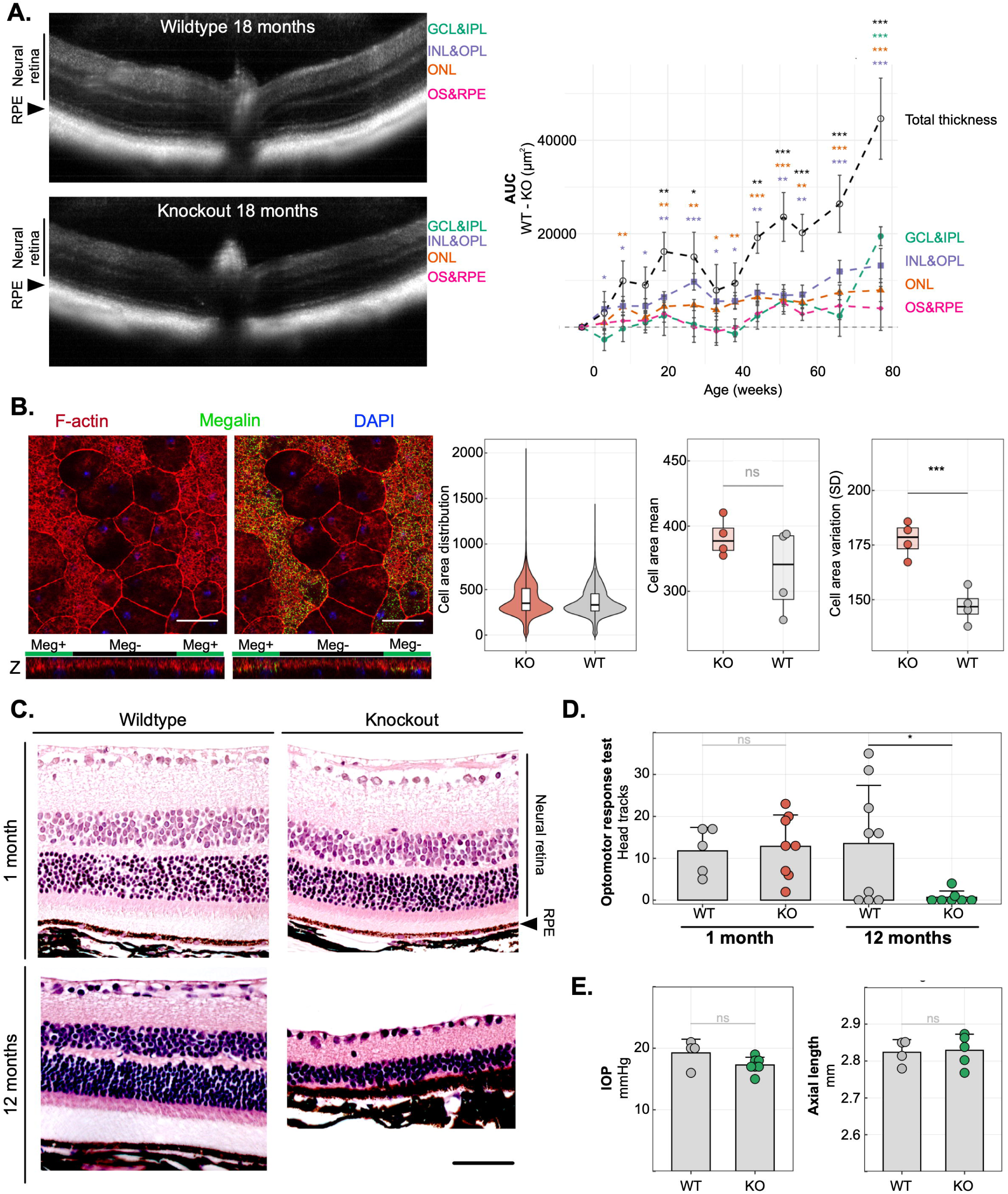
Megalin ablation causes retinal degeneration in mice. **A)** Representative Optical Coherence Tomography scans of retinas from WT and megalin KO mouse at 18 months and quantitation of the thickness of each retinal layer over time presented as difference in Area Under the Curve (AUC) between WT and KO. **B)** Representative immunofluorescence of RPE flatmount from a megalin KO mouse stained for megalin (green), actin (red) and DAPI (blue). RPE cell heterogeneity, measured as cell area distribution and mean of cell areal was comparable between megalin-expression and megalin-deficient cells in megalin-KO mice. Cell area variation was increased in megalin-deficient cells compared to megalin-expressing. Scale bar: 20 µm. **C)** Hematoxylin/eosin staining at 1 and 12 months in megalin KO mice and WT controls. Scale bar: 50 µm. **D)** Optomotor response test in megalin-KO mice 1 month and 12 months after KO. **E)** Intraocular pressure (IOP) and axial length measurements in WT and KO mice after 12 months.

Although overall retinal architecture remained largely preserved after three months, RPE flatmounts of megalin-KO mice revealed pronounced morphological changes in megalin-deficient cells at this time. Megalin-deficient RPE cells appeared enlarged and irregular, with loss of the characteristic hexagonal morphology, whereas neighboring megalin-positive cells appeared compressed (Fig. 2B and Supplementary Fig. S1). While mean RPE cell area was not significantly altered between WT and KO animals, KO RPE exhibited markedly increased variability in cell size, indicating disrupted epithelial organization and early cellular stress.

Consistent with the OCT findings, histological analysis confirmed preserved retinal morphology at one month, whereas pronounced thinning of the outer retina was observed in some animals after one year (Fig. 2C). A subset of the included animals displayed a milder phenotype (Supplementary Fig. S2), indicating variability in disease progression. In accordance with histological findings, visual function assessed by optomotor response test remained intact four weeks after induction of KO but was impaired at 12 months, indicating progressive retinal dysfunction (Fig. 2D). In contrast to mouse models with constitutive megalin-KO[14, 16], axial length and intraocular pressure were unchanged between groups after 12 months (Fig. 2E and Supplementary Fig. S3), suggesting that degeneration was not secondary to global ocular alterations.

### Alterations in autophagy-related processes related to POS disc trafficking

To investigate the overall changes in megalin-deficient RPE cells, mass spectrometry-based proteomic profiling on isolated mouse RPE cells from KO and WT mice at 3 months was performed. This revealed downregulation of proteins primarily involved in ion channel transport and autophagy, whereas pathways associated with cellular health were upregulated including DNA double strand repair and epigenetic regulation (Fig. 3A). To ensure that these changes were not mouse specific and could be extrapolated to humans, we also performed proteomics on human iPSC-RPE cells comparing control (Ct) cells to KD cells using aptamer-based proteomics (Fig. S4). Likely due to the lower KD efficiency in the cell model, log-fold changes (logFC) were generally lower in iPSC-RPE cells compared to mouse RPE and we therefore compared proteomes based on rank-percentile rather than raw log values. This revealed a significant overlap between the two proteomes visualized as green dots representing proteins that are consistently downregulated in both datasets and red dots representing proteins that are consistently upregulated (Fig. 3B). Since selective autophagy was one of the most significantly downregulated pathways in mice, and the autophagy machinery is used to traffic POS discs in the RPE, we used Gene set enrichment analysis (GSEA) to test if this pathway was also reduced in KD iPSC-RPE cells. This revealed a significant reduction of autophagy-related proteins in megalin-KD cells (NES = −1.82, p = 0.01) (Fig. 3C), indicating a coordinated reduction across the pathway in both models. When the two proteomic datasets were combined, autophagy was the most significantly reduced pathway following megalin-ablation (Fig. 3D).

**Figure 3.**
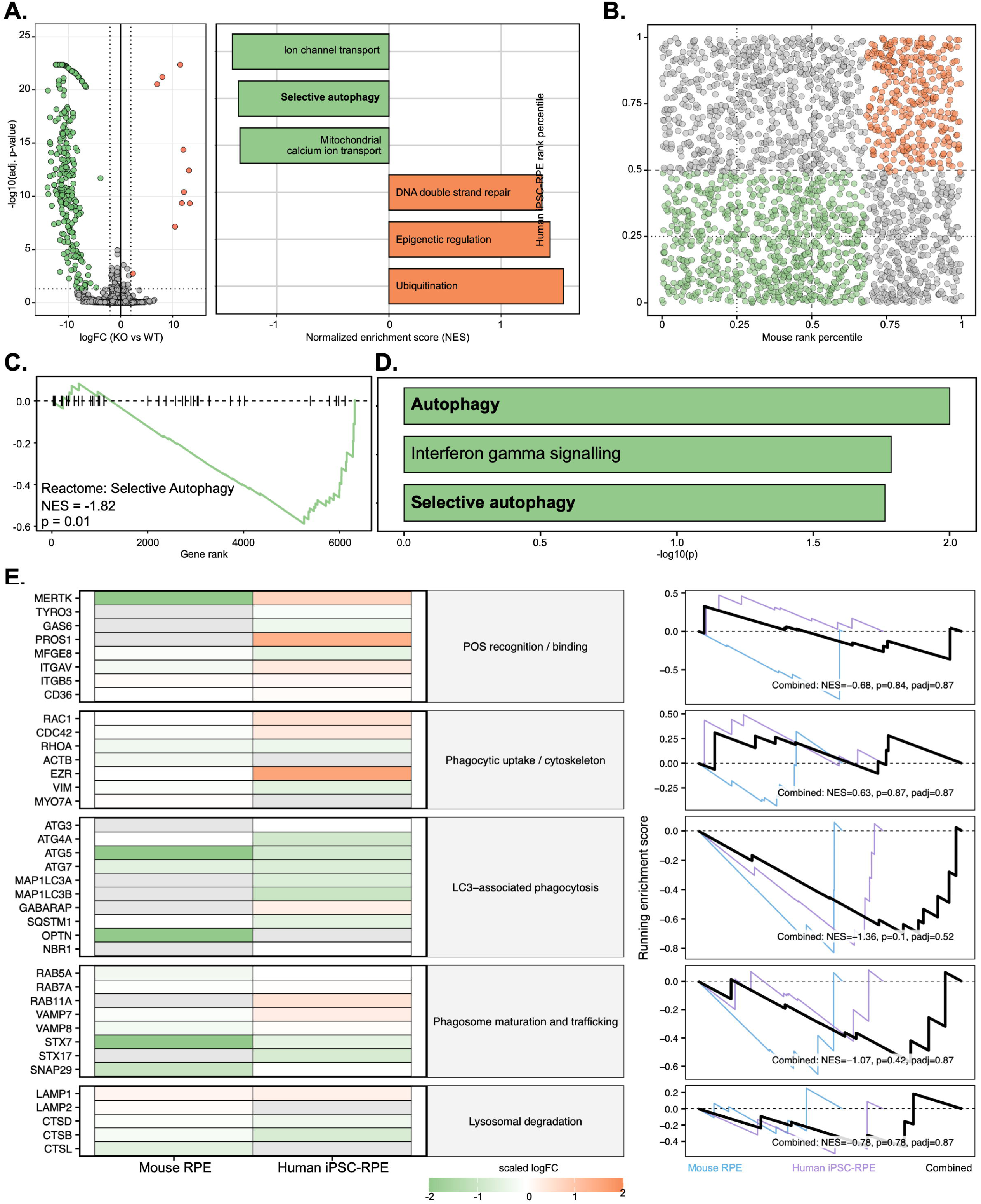
Proteomics reveal changes in autophagy in isolated mice RPE and human iPSC-RPE cells. **A)** The proteomic signature of isolated RPE cells from megalin-KO mice compared to WT controls. Volcano plot shows up- and downregulated proteins (p<0.05, logFC >2). Unsupervised pathway analysis using the Reactome pathway database showing the most significantly up-and downregulated pathways. **B)** Rank-rank concordance plot comparing mouse RPE proteome to human iPSC-RPE proteome. Green dots represent proteins that are consistently downregulated in both datasets and red dots represent proteins that are consistently upregulated. **C)** Gene Set Enrichment Analysis (GSEA) of megalin KD cells compared to controls using the Reactome autophagy. **D)** Unsupervised Reactome pathway analysis of combined mouse RPE cells and human iPSC-RPE cells. **E)** Heatmap of proteins involved in POS trafficking and degradation in RPE. LogFC are scaled after SD of each dataset. GSEA of each sub-pathway are shown for mouse, human and combined. Green is downregulated, orange is upregulated.

To localize the origin of this signal within the pathway, autophagy-related proteins were grouped into functional sub-pathways and logFC were scaled based on standard deviation within each dataset. GSEA was performed for each sub-pathway for human iPSC-RPE, mouse RPE cells as well as the combined proteome. Although none of these analyses reached significance, proteins involved in LAP showed consistent downregulation across most proteins including ATG5 in both datasets (Fig. 3E). Since this is a major pathway for POS disc trafficking in the RPE, this may indicate that megalin is a potential regulator of LAP.

### Megalin deficiency impairs rhodopsin degradation

Given the coordinated changes across the autophagy machinery and LAP as well as the established role of megalin in endocytosis in other tissues, we hypothesized that megalin contributes to the handling of POS discs in the RPE. To test this, iPSC-derived RPE cells were exposed to FITC-labelled POS discs isolated from bovine retinas, and intracellular processing was monitored over time (Fig. 4A). Control and megalin-KD cells showed comparable uptake after 2 hours, as determined by similar rhodopsin levels, indicating that initial POS disc phagocytosis was not affected in this short *in vitro* assay. However, western blot analysis revealed delayed rhodopsin degradation in megalin-KD cells compared to controls (Fig. 4B).

**Figure 4:**
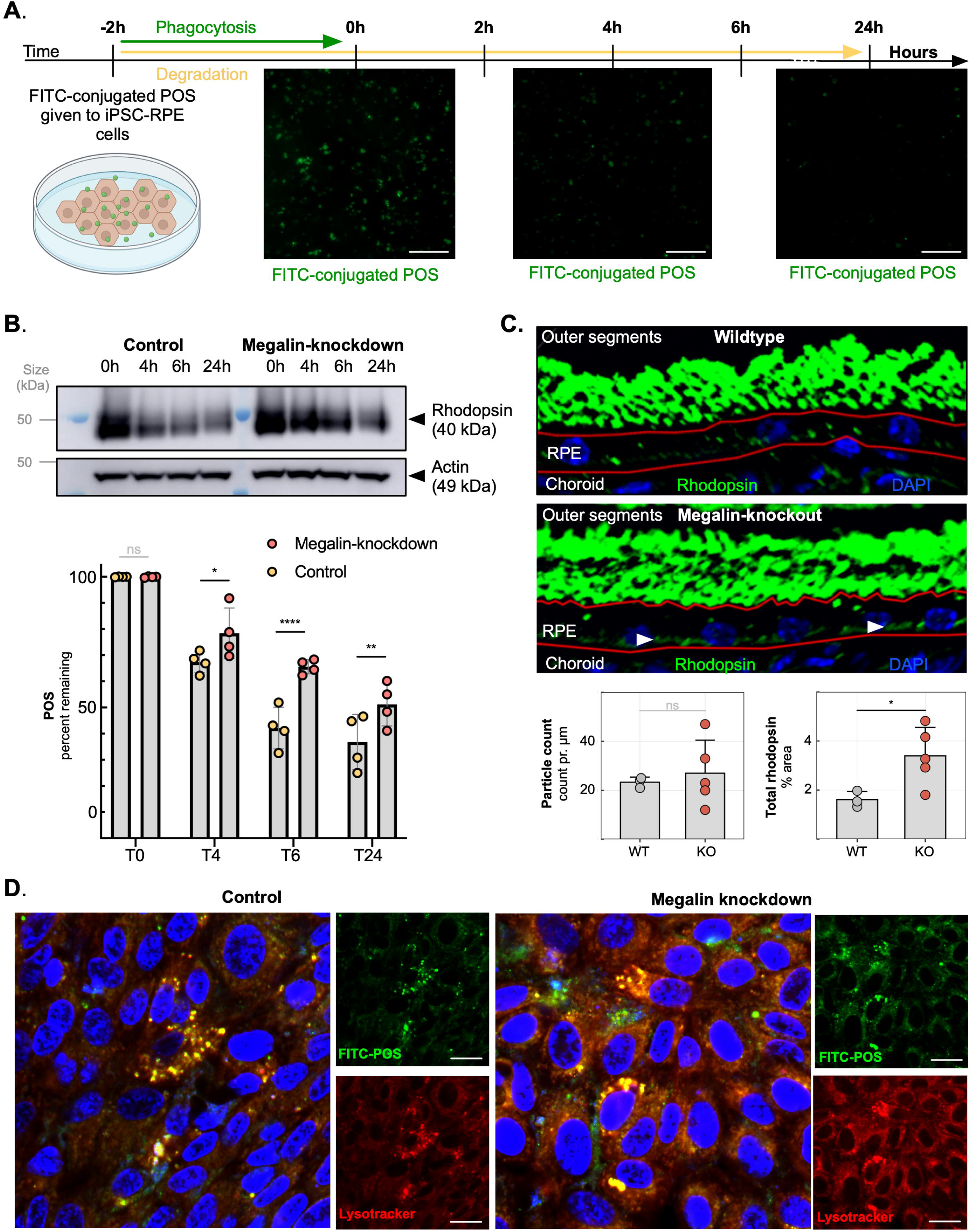
Photoreceptor outer segment turnover is slower in megalin-knockdown RPE. **A)** Immunofluorescence of iPSC-RPE cells fed with FITC-conjugated bovine outer segments (green) for two hours (pulse) and subsequent degradation (chase). Scale bar is 100 µm. **B)** Representative western blotting of rhodopsin in control and megalin KD cells immediately after POS pulse and up to 24 hours after. **C)** Representative immunohistochemistry of rhodopsin (green) and DAPI (blue) on paraffin-sections of mice retinas for quantification of particle counts and total rhodopsin content. **D)** Immunocytochemistry performed two hours after incubation of iPSC-RPE with FITC-POS (green), lysotracker (red) and DAPI (blue). Scale bar is 20 µm.

To determine whether POS disc processing was changed *in vivo*, rhodopsin accumulation was assessed in megalin-KO mice. While quantification of rhodopsin-positive particles in the RPE showed a non-significant trend toward increased particle counts in megalin-KO mice one month after megalin ablation, megalin-KO mice exhibited increased overall rhodopsin-positive area within the RPE. This was accompanied by a diffuse basal rhodopsin staining pattern that was not observed in control animals (Fig. 4C; white arrows), suggesting accumulation of incompletely degraded POS disc material.

To examine whether megalin deficiency affects phagosome maturation and lysosomal trafficking of POS discs in megalin-KD iPSC-RPE cells, we performed immunocytochemistry of FITC-conjugated POS discs together with LysoTracker. This revealed that, after 2 hours, most POS discs localized to lysosomes in control cells, whereas a substantial fraction of POS discs remained outside lysosomal compartments in megalin-KD cells (Fig. 4D) suggesting impaired phagosome maturation causing ineffective fusion with lysosomes.

### Megalin deficiency leads to accumulation of pro-cathepsin D without loss of lysosomal activity

To determine whether these alterations were caused by lysosomal dysfunction, RPE/choroid cross-sections were stained for rhodopsin and cathepsin D, which is the dominant lysosomal protease required to achieve efficient rhodopsin degradation. Only few rhodopsin-positive structures co-localized with cathepsin D in both WT and KO mice. However, KO mice exhibited increased cathepsin D particle density compared with WT animals (mixed-effects model, p = 0.028; Welch’s t-test, p = 0.035; Cohen’s d = 2.17) (Fig. 5A), indicating cathepsin D accumulation in the absence of megalin.

**Figure 5:**
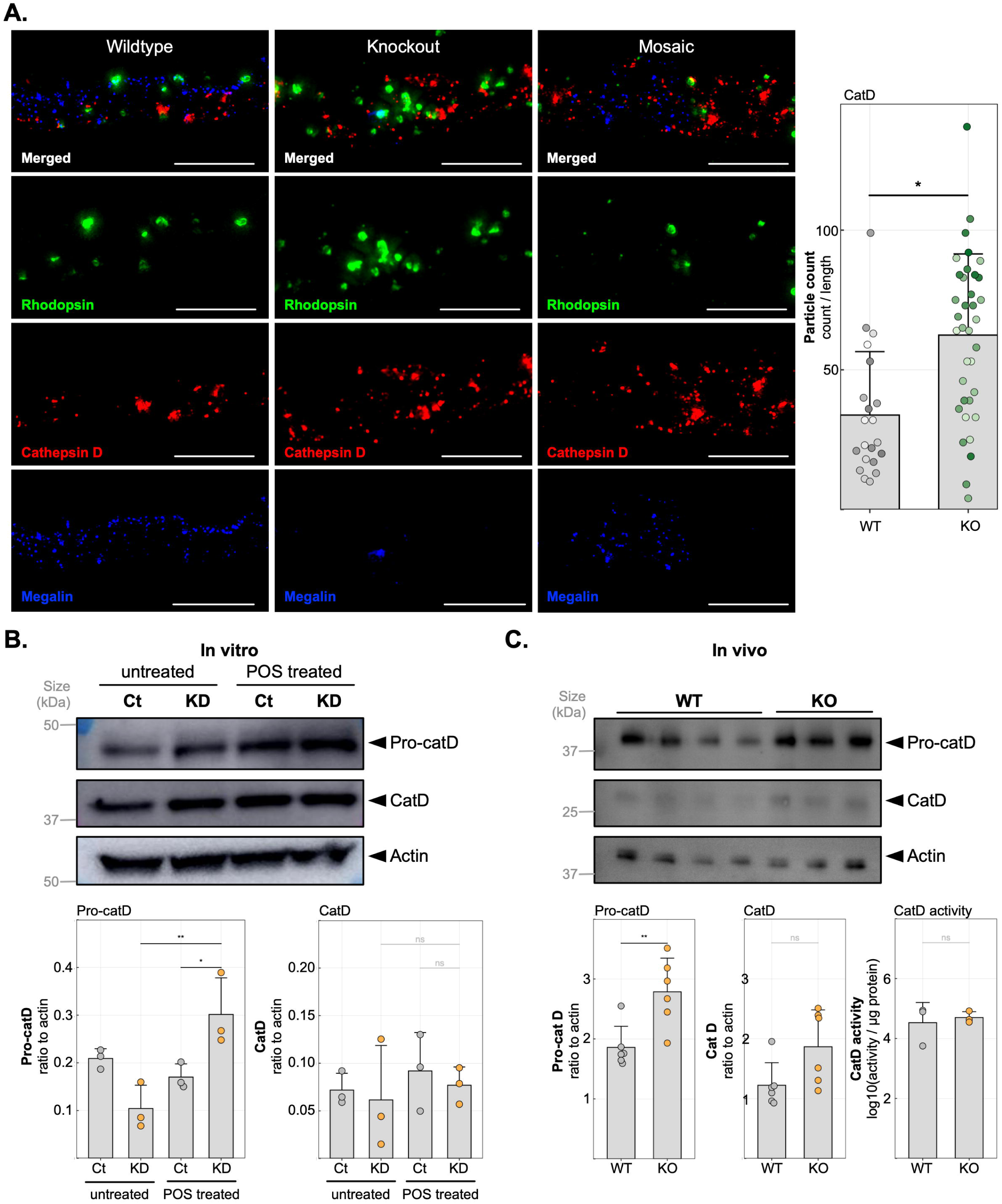
Lysosomal protease maturation is disrupted upon megalin ablation. **A)** Immunohistochemistry of paraffin-embedded RPE sections of megalin KO and WT mice as well as an example of mosaicism stained for rhodopsin (green), cathepsin D (red) and megalin (blue). Cathepsin D particle counts were quantified and compared between megalin KO and WT. Scale bar: 10 µm. **B)** Quantitative western blotting of cathepsin D showing both immature pro-cathepsin D and mature cathepsin D bands in iPSC-RPE both under basal conditions and fed with POS. **C)** Quantitative western blotting of cathepsin showing both immature pro-cathepsin D and mature cathepsin D in isolated mouse RPE cells. All protein levels were compared to actin. Cathepsin D fluorometric activity assay performed on isolated mouse RPE cells.

Western blot analysis of pro-cathepsin D and cathepsin D *in vitro* and *in vivo* revealed that the increased cathepsin D signal observed in megalin-KO RPE was primarily driven by accumulation of pro-cathepsin D rather than the mature protease. *In vitro*, pro-cathepsin D accumulation in megalin-KD RPE was observed specifically following POS disc exposure, linking this phenotype to impaired POS disc processing (Fig. 5B). *In vivo,* where POS disc exposure is constant under physiological conditions, pro-cathepsin D accumulation was also observed. To determine whether lysosomal proteolytic capacity was impaired, cathepsin D activity was assessed *in vivo* using an artificial cleavage substrate GKPILFFRLK(Dnp)-D-R-NH2). No difference in cathepsin D activity was detected between WT and KO animals (Fig. 5C), indicating that accumulation of POS discs was not caused by a mature cathepsin D-associated lysosomal deficit.

## Discussion

Using inducible post-developmental KO models in mice and human iPSC-derived RPE, we identify megalin as an important regulator of mature RPE homeostasis and retinal integrity. In contrast to developmental knockout models, inducing megalin loss in the mature retina did not alter axial length or intraocular pressure, but instead led to morphological abnormalities in the RPE, progressive retinal degeneration, and visual decline. These findings support a continued requirement for megalin in the adult retina, independent of its crucial developmental roles. Mechanistically, our data suggests that megalin deficiency impairs POS disc processing due to disrupted LC3-associated phagosome maturation and trafficking rather than a primary lysosomal dysfunction. This disruption may potentially delay fusion with lysosomes resulting in delayed rhodopsin degradation, accumulation of incompletely processed POS disc material, and elevated levels of pro-cathepsin D.

Megalin deficiency caused morphological changes in both the RPE and the neural retina. In the RPE, megalin-KO cells exhibited larger variability in cell size compared to WT cells. Proteomic analyses revealed changes in pathways like TGF-beta signaling, transcription and cell death. Together, these results indicate a decline in overall cellular health following megalin depletion consistent with studies of dedifferentiation and epithelial to mesenchymal transition (EMT)[23–25]. These changes are later echoed in the inner retina where progressive retinal thinning is observed. This pattern of RPE dysfunction followed by slowly progressive retinal degeneration is similar to that observed in RPE65-KO mice[26].

Proteomic analysis of isolated mouse RPE cells and iPSC-RPE revealed coordinated changes in the autophagy machinery, which is used to degrade ingested POS discs in the RPE through the process of LAP. The RPE is exposed to a continuous and unusually high degradative burden from daily POS disc clearance and utilizes the autophagy machinery to effectively degrade POS discs. Negative enrichment of autophagy proteins was primarily driven by reduced ATG5, MAP1LC3A/B, ATG4A, and ATG7, all of which are required for LC3 processing and lipidation. Because LC3 conjugation is critical for LAP and phagosome maturation in the RPE, these data suggest that megalin deficiency disrupts coupling of POS disc phagosomes to downstream degradative pathways rather than causing a broad collapse of lysosomal function.

We further studied how megalin deficiency affected POS disc uptake and subsequent rhodopsin clearance both *in vitro* and *in vivo*. POS disc engulfment was unimpaired *in vitro*, indicating that megalin is not required for the initial phagocytic uptake of POS discs under these conditions. However, rhodopsin degradation was delayed in megalin-deficient iPSC-RPE, and POS discs showed reduced co-localization with lysosomes, consistent with impaired progression from phagosome formation to degradative fusion. When investigating rhodopsin turnover *in vivo*, rhodopsin particle counts were comparable between genotypes, but larger rhodopsin-positive aggregates accumulated toward the basal side of the RPE in KO retinas. Although POS disc engulfment was unimpaired *in vitro*, we cannot exclude that a defect in the system *in vivo* will feedback into the phagocytic activity. Downregulation of MERTK in the mouse model may indicate such a defect and explain both the unchanged rhodopsin particle counts and the appearance of diffuse basal rhodopsin in KO cells. These experiments point to a defect in post-phagocytic processing, and megalin appears to be required for efficient trafficking and degradation of cargo. Since even modest defects in POS disc clearance may have major consequences in the RPE due to its continuous and exceptionally high phagocytic burden[10], this may explain how a relatively subtle cellular defect can translate into progressive retinal degeneration over time.

We next asked whether the impaired POS disc degradation reflected defective lysosomal enzyme maturation or a broader inability to deliver POS disc-containing phagosomes to functional lysosomes. Levels of mature cathepsin D were comparable between genotypes, whereas pro-cathepsin D levels accumulated in KD-cells specifically upon POS challenge, but not under basal culture conditions. This context-dependent accumulation suggests that the defect is not a consequence of lacking constitutive megalin-mediated endocytosis of cathepsins but rather it emerges from the POS load itself and the demand it places on the degradative machinery. Consistent with intact lysosomal biogenesis, mature cathepsin D abundance and activity were comparable between genotypes *in vivo*. Notably, the cathepsin D profile observed in cultured cells was recapitulated *in vivo*, where POS disc degradation is a constant process. The accumulation of pro-cathepsin D could thus be a result of not using enough of the mature protease or a failed stress response of attempting to increase proteolytic capacity. A similar phenotype has previously been described in chloroquine-treated RPE cells where accumulation of pro-cathepsin D was also observed[10]. Chloroquine works as an inhibitor of autophagic flux and decreases autophagosome-lysosome fusion[27]. Together, these observations indicate that the degradative deficit in megalin-deficient cells is not explained by altered lysosomal biogenesis but rather reflects an inability to appropriately scale phagosome-lysosome fusion to the POS disc load.

Overall, our results establish megalin as an important player in phagosome-to-lysosome trafficking in parallel with the established role of megalin in renal handling of lysosomal proteins[28] and support a broader function for megalin in sustaining phagosome maturation, here specifically in the context of POS disc clearance by the RPE. Our findings support a model in which megalin facilitates coupling of POS disc-containing phagosomes to downstream lysosomal degradation. Reduced lysosomal co-localization of POS discs suggests inefficient progression from phagosome formation to degradative fusion. Megalin may therefore function as an organizer of intracellular trafficking rather than simply as a surface uptake receptor. These findings suggest that reduced megalin function may lower the capacity of the RPE to withstand chronic degradative stress. In other tissues, megalin-expression is known to decline with age[29, 30]. Such vulnerability may be relevant in retinal disorders characterized by lysosomal burden and incomplete cargo clearance such as AMD[31, 32]. Our data therefore positions megalin-dependent pathways as potential modifiers of retinal degenerative disease in accordance with previous work linking megalin to high myopia, retinal dystrophy, and lysosomal dysfunction[14]. The results presented extend these observations further by demonstrating that megalin remains functionally important in the adult RPE. An important remaining question is whether megalin acts directly in LAP-related membrane dynamics or more broadly in phagosome-lysosome organization. Future studies will be needed to define whether the observed phenotype reflects a primary defect in LAP maturation, fusion with lysosomes, or both. Future studies will also need identify which vesicles megalin localizes to.

The concordance between the mouse and human cell models is a major strength of the study. Limitations include the partial knockdown achieved in iPSC-RPE and the fact that our data do not yet resolve whether megalin acts directly in LAP-associated trafficking or more broadly by regulating phagosome-lysosome fusion. While the mosaic nature of the mouse model does cause the knockout degree to be slightly lower, it also allows us to study neighboring megalin-positive and megalin-negative cells within a single animal, which contributes important information to the analysis.

Collectively, we identify megalin as a key regulator of lysosomal health and POS disc processing in the RPE. Megalin deficiency disrupts phagosome maturation and trafficking, is associated with reduced expression of the LC3 conjugation machinery and altered cathepsin D processing and ultimately compromises retinal integrity *in vivo*. These findings position megalin as a candidate regulator of RPE resilience and a potential point of vulnerability in retinal degenerative disease.

## Methods

### iPSC Culture and Differentiation

The PGP-1 WT P38 line was acquired from Synthego (Menlo Park, CA). Since the cell lines carried heterozygous high-risk AMD variants at the ARMS2/HTRA1 site (ARMS2 G/T, HTRA1 G/A), these were CRISPR-edited back to wildtype alleles (ARMS2 G/G, HTRA1 G/G) using Synthego’s industrialized CRISPR platform. All iPSCs were maintained in mTeSR™ Plus (STEMCELL, cat. 100-0276) on Matrigel-coated plates (corning, product number 354234) and passaged every 3-7 days. iPSC cultures (day 0) were cultured for 7□days in neural induction medium which consisted of 50% DMEM/F12□+□GlutaMAX (GIBCO), 50% neurobasal medium (GIBCO), 0.5x B27 supplement (GIBCO), 0.5x N2 supplement (GIBCO), 55□μM 2-mercaptoethanol (GIBCO), and 1x L-glutamine (GIBCO). Medium was changed every 3□days. At day 7, cells were washed twice in PBS and cultured in RPE medium with□50□ng/ml recombinant human Activin A (PeproTech) until first signs of pigmentation were visible with the naked eye (normally at days 18–21). RPE medium is formulated as follows: DMEM, high glucose, GlutaMAX™ Supplement (GIBCO), 10% KnockOut Serum Replacement (GIBCO), 1x L-glutamine (GIBCO), 1x MEM Non-essential Amino Acid Solution (Sigma-Aldrich), 55□μM 2-mercaptoethanol (GIBCO), 1x penicillin-streptomycin (GIBCO). Medium was changed every 3□days. As soon as pigmentation started, cells were cultured in RPE medium without Activin A and the culture was carried on until at least 35–40% of the culture plate surface showed pigmentation (normally around day 35).

### shRNA-mediated LRP2 knockdown

shRNA in a lentiviral particle was obtained from OriGene (Cat. #TL311679V). Four shRNA constructs were screened, and the most efficient was used for our experiments. RPE cells were allowed to differentiate for 3 weeks before knockdown was achieved by treating the cells with 25 MOI of either scrambled or LRP2-shRNA for 16 hours along with 0.8 µg/ml polybrene. For mRNA measurements, cells were left for additional 24 hours in complete media. For proteins measurements, cells were left for 72 hours after knockdown before experiments were begun.

### Semi quantitative RT-PCR

Total RNA from the differentiated iPSC-RPE (∼10^6^ cells) was purified by using the RNeasy micro kit (Qiagen, Cat. 74004) according to the manufacturer’s instruction. Total 1 µg of RNA was reversely transcribed using Omniscript RT Kit (Qiagen, Cat. ID: 205111). Using 1 µL of cDNA, LRP2 fragment was amplified using Platinum II Taq Hot-Start DNA Polymerase (Invitrogen, Cat. 14966001) and forward and reverse primers (LRP2 Forward – AGCCTCTGGAGTTGGACAGA, Reverse - ACAGTGCGGTTAGACCCATC; 125 nM). PCR method is as follows: 94°C 5 min; 94°C 30 S, 58°C 30S, 72°C 45 S with 35 cycles; 72°C 10 min; product size: 189bp. Relative amount of LRP2 mRNA was compared by measuring the relative band intensity of PCR amplicons by electrophoresis (Bio-Rad CFX96 Touch, Bio-Rad Laboratories, CA) on 1% agarose gel. Control beta-actin (Forward -GGACTTCGAGCAAGAGATGG, Reverse - AGCACTGTGTTGGCGTACAG) was amplified with PCR method described as followed: 94°C 5 min; 94°C 30 S, 58°C 30S, 72°C 45 S with 35 cycles; 72°C 10 min; product size: 214 bp.

### Western blotting

#### In vitro

Cells were lysed using RIPA buffer and protein was extracted and centrifuged at 300g for 5 minutes. Total protein concentration was measured using the BCA assay (ThermoFischer cat#A55864) and normalized to 5ng. Samples were loaded onto a 4-12% Bis-Tris gel with a high molecular weight marker and separated by 220V for 1 hour and 15 minutes. Gels were transferred to a PVDF membrane using the iBlot3 (ThermoFischer, USA) system for 8 minutes before blocking in 5% milk for 1 hour. Primary antibody (sheep-anti-megalin) has been described and characterized previously[33]. Incubation was performed at 4°C overnight and blots were washed in TBS-T through SNAP i.d.® 2.0 Protein Detection System (Bio-Rad Laboratories, CA). Secondary antibody binding was performed for 30 minutes at room temperature before additional washing in TBS-T. Blots were visualized using the SuperSignal West Dura Extended Duration Substrate (Thermo Scientific™, MA) was added to the membranes to develop immunochemical signal, and the chemiluminescent profile was visualized with Amersham Imager 600 (GE Healthcare Life Sciences, IL).

#### In Vivo

RPE cells were isolated by making a circular cut around the iris and removing the lens to expose the eyecup. Trypsin digestion was performed for 1 hour at 37°C and RPE cells were rinsed from the internal surface of the retina. Samples were loaded onto a 16% Tris-Glycine gel or 3-8% Tris-Acetate gel (NuPage, Invitrogen) with a standard molecular weight marker (Invitrogen) and separated by 42 minutes of electrophoresis at 225V. The proteins were then transferred to a PVDF fluorescence membrane or a PVDF membrane for ECL (Millipore) using the Invitrogen iBlot System for 3-7 minutes depending on protein size, at 20V. Blots were blocked using 5% skim milk (Millipore Sigma #1.1563.0500 in PBS) for one hour. Blots were then incubated with primary antibodies at 4°C overnight and washed in PBS-Tween to remove unbound antibodies. Blots were then incubated with or peroxidase-conjugated secondary antibodies for 1 hour. The membranes were again washed in PBS-Tween to remove unbound antibodies. Membranes were visualized using Immobilon Western Chemiluminescent HPR Substrate (Millipore) and the Image Quant Las 4000. Protein content was compared to content of actin to ensure correct analysis.

### POS uptake and degradation

Bovine photoreceptor outer segments (POS) were kindly provided by Stephen H. Tsang, Columbia University, and used for RPE phagocytosis/degradation assays as previously described in studies from the Tsang laboratory[34] and according to established POS purification protocols[35]. Experiments were performed in 12-well plates with 1 x 10^6^ POS pr. ml of media. For immunocytochemistry, POS were conjugated to an FITC molecule (Fluorescein 5(6)-isothiocyanate, Sigma, #46950) for 1 hour in the dark following a previously published protocol[36]. Lysotracker staining was achieved by incubating cells with LysoTracker red DND-99 (Thermo Fisher Scientific, L7528) at a final concentration of 100 nM in culture medium.

### Immunocytochemistry

Cells were fixed in 4% PFA, permeabilized with 0.2% Triton-X, and blocked in 5% BSA in PBS. Primary antibody incubation with LRP2 (abcam, cat. ab76969), Occludin (Invitrogen, cat. 33-1500), ZO-1 (Alexa Fluor 488 conjugate, Invitrogen, cat. MA3-39100-A488), CRALBP (abcam, cat. ab15051) and RPE65 (sigma-aldrich, cat. MAB5428) to characterize the RPE monolayer was performed overnight at 4 °C. Except for ZO-1, the cells were incubated with secondary antibody (GeneTex, CA, Cat # GTX213111-04 or −05, goat anti-Mouse IgG-DyLight594 conjugate or DyLight488 conjugate, 1/1000). After three times of wash with 0.1% TritonX100 in PBS, the cells were covered with VECTASHIELD® Antifade Mounting Medium with DAPI (Vector Laboratories, CA, Cat # H-1200-10). Images were obtained with Zeiss Laser confocal microscope LSM 880.

### Animal model

Meg^lox/lox^ mice were crossed with tamoxifen-inducible UBC (ubiquitin C)-cre/ERT2 (mutant form of the estrogen recetor) transgenic mice (The Jackson Laboratory, Bar Harbor, ME) to produce Cre^+^, meg^lox/lox^ mice. All mice were on a 129SvE-F Tac IK background. At age 8-12 weeks, female megalin KO (Meg^lox/lox^; Cre^+^) (KO) and littermate control mice (Meg^lox/lox^; Cre^-^) (WT) were induced by i.p. injection of tamoxifen (Sigma-Aldrich, St. Louis, MO) at a dose of 100 mg/kg body weight for 5 consecutive days. The ubiquitin C promotor has a widespread activity and knockout of megalin is expected in all tissues. Decrease of megalin was checked either by western blotting of isolated RPE cells or by immunohistochemistry of the retina for megalin. During some experiments, all material was used and KO degree was extrapolated from the kidney. Animals were anesthetized with isoflurane for tamoxifen injections and euthanasia. For all other procedures, anesthesia was induced by intraperitoneal injection of ketamine (60 mg/kg) and medetomidine hydrochloride (1 mg/kg). Anesthesia was reversed by intraperitoneal injection of atipamezole hydrochloride (1 mg/kg). For optical coherence tomography (OCT), pupils were dilated using tropicamide eye drops (Mydriacyl, 0.5–1%), and ViscoTears eye gel was applied to prevent corneal dehydration and irritation during imaging. Mouse breeding and experiments were carried out in the certified animal facility at Aarhus University according to the Danish Animal Experiments Inspectorate and under permission no 2023-15-0201-01585.

### Proteomics

#### In vitro

Cells were cultured in 12-well plates for 3 weeks in complete media. Cells were washed with PBS and lysed in 150 µL M-PER buffer containing protease inhibitor according to the manufacturer’s instructions. Samples were diluted to a total protein concentration of 200 µg/mL in PBS before shipment to Somalogic for aptamer-based proteomics. Bioinformatic analyses were performed as previously described[37–39] using R Studio (version 2022.02.0+443, R version 4.1.2) with the following packages: limma, ggplot2, clusterProfiler and reactomePA,

#### In vivo

Mass spectrometry (MS) proteomics were performed on isolated mouse RPE cells from mice aged 3 months. 7 WT and 7 KO mice were included. Peptides were prepared using the filter-aided sample preparation (FASP) method. Cells were dissolved in PBS containing 1% SDS with protease and phosphatase inhibitors. After sonication, proteins from each animal were loaded to a Vivacon-30 kDa spin column and centrifuged at 16,000 × g to remove all liquids. The columns were washed three times with a UA buffer containing 8 M urea and 100 mM triethylammonium bicarbonate. Proteins were then reduced with 50 mM dithiothreitol for 1 hour at 37□°C, followed by alkylation with 50 mM iodoacetamide for 20 minutes at room temperature in the dark. After each step, centrifugation was performed to remove all liquids. Samples were then once again washed in UA buffer before adding 5 µl of Lys-C solution at an enzyme-to-protein ratio of 1:50, dissolved in UA buffer. The mixture was incubated at 37□°C for 2 hours, and 50 µl of trypsin solution (enzyme-to-protein ratio of 1:25, dissolved in 100 mM triethylammonium bicarbonate) was added. Digestion was performed overnight at 37□°C. Next morning, 4uL of 10% TFA was added to the samples which were then centrifuged to collect the peptides. Desalting was performed using C18 desalting tips. After desalting, samples were lyophilized in a SpeedVac until dry. The samples were stored in −80°C until LC-MS analysis.

LC-MS analysis was performed using Vanquish Neo HPLC coupled with an Orbitrap Ascend MS and a FAIMS Pro Duo interface. A 30-minute gradient with an active separation window of 24.5 minutes of 5-40% of B (80% ACN and 0.1% FA) was used to separate the peptides on a 15-cm long Ion Opticks Aurora column operated under 50°C. MS analysis was done in DIA (Data Independent Acquisition) mode, with FAIMS CV of −50. Parameters for MS1 scans include scan rage of 380-980, Orbitrap resolution of 120K, maximum ion injection time of 251ms, normalized AGC of 250%. Parameters for MS2 scans include DIA window size of 40Da, Orbitrap resolution of 120K, maximum ion injection time of 251ms, normalized AGC of 1000%, collision energy of 30%, number of scan event of 15, loop count of 5 (one MS1 scan inserted between every 5 MS2 scans), and loop time of 3 seconds.

The MS raw data was analyzed by direct DIA mode in Spectronaut (v20) against a mouse uniprot database downloaded on March 20, 2026. Peptide and protein false discovery rates were set at 1%. All parameters were default except the quantification level set to MS1. Identified proteins and the quantification matrix were exported to a csv file for downstream data analysis. Differential protein abundance was visualized by volcano plot with significantly altered proteins defined by adjusted p-value < 0.05 and absolute log2 fold change > 2.

Reactome gene set enrichment analysis was performed using ranked protein lists ordered by log2 fold change. Enrichment was assessed using normalized enrichment score (NES), and adjusted p-value. To compare mouse and human datasets, proteins detected in both datasets were ranked by log2 fold change within each dataset and plotted against each other as percentile ranks. Proteins were classified according to concordant or discordant direction of change between mouse and human datasets.

Shared down-ranked proteins were defined as proteins ranking within the most downregulated quartile in both datasets. Reactome overrepresentation analysis was then performed using R. Proteins contributing to selected enriched pathways were extracted for visualization.

To assess POS-related cargo handling log2 fold changes were scaled within each dataset by the dataset-specific standard deviation to preserve direction while allowing comparison across platforms. Proteins absent from one dataset were shown in grey. Custom enrichment curves were generated for each curated category in mouse, human, and combined ranked lists, with combined rankings based on the mean scaled log2 fold change across datasets.

### Optomotor response test

Visual function was assessed using an optomotor response test at 1 month and 1 year after induction of megalin knockout. Mice were placed individually on a platform in the center of a rotating optomotor drum measuring 30 cm in diameter and displaying a black-and-white vertical striated pattern with a stripe width of 3 mm, corresponding to 1.15°. The drum was rotated at 4 rpm. Each mouse was allowed to acclimatize on the platform for 1 min before the drum rotation was initiated.

Behavior was recorded from above for 2 min using a Polaroid XS100 camera mounted centrally above the platform. Each mouse was tested three times. Recordings were blinded and randomized by a person not involved in the optomotor response testing before analysis. Head tracking was defined as a rotational head movement in the same direction and at the same speed as the rotating drum for at least 2 s. The total number of head-tracking events across the three trials was used for comparison between genotypes.

### Intraocular pressure measurement

Intraocular pressure was measured 1 year after induction to assess whether retinal degeneration was associated with altered ocular pressure. Measurements were performed using the TonoLab rebound tonometer (TonoVet, Finland). Mice were anesthetized with ketamine/domitor, and both eyes were measured in each animal. For each eye, 3 readings were obtained, and the mean value was used as the eye-level intraocular pressure. The mean of both eyes was then used as the animal-level value for downstream statistical analysis. Measurements were performed blinded to genotype. Intraocular pressure was compared between megalin-KO and littermate control mice using student’s t-test.

### Axial length measurements

Axial length was measured at 67 weeks using a Vevo F2 LAZR-X ultrasound imaging system equipped with a UHF46x transducer (20–46 MHz, 50 µm axial resolution; FUJIFILM VisualSonics). Mice were anesthetized with ketamine, and both eyes were imaged in each animal. Axial length was measured directly in the Vevo software on B-mode ultrasound images as the distance from the anterior corneal surface to the posterior pole of the eye along the anteroposterior axis (Fig. S3). One measurement was obtained per eye, and the mean of both eyes was used as the animal-level axial length for downstream analysis. Measurements were performed blinded to genotype. Axial length was compared between groups as a control measurement to assess whether retinal degeneration was associated with gross differences in eye size.

### Histology and paraffin sectioning

Eyes were fixed in 4% PFA, embedded in paraffin using standard procedures, and sectioned at 5 µm thickness. Sections were mounted on SuperFrost Plus slides and dried for at least 60 min at 60°C. Before staining, sections were deparaffinized in xylene overnight and rehydrated through graded ethanol solutions of 99%, 96%, and 70%, followed by several washes in distilled water. Sections were used for hematoxylin and eosin (HE)-staining or immunofluorescence staining.

For HE staining, sections were incubated in Mayer’s hematoxylin for 5 min and washed in running tap water for 5 min. Sections were then incubated in eosin for 2 min, dehydrated through 96% and 99% ethanol, cleared in xylene, and mounted with Eukitt mounting medium.

For immunofluorescence, paraffin sections were deparaffinized, rehydrated, and subjected to antigen retrieval by boiling in TEG buffer containing 0.01 M Tris-base and 0.0005 M EGTA in a microwave for 10 min. Sections were allowed to cool for 30 min before incubation in 50 mM NH_₄_Cl in PBS for 30 min to block free aldehydes. Sections were blocked in buffer containing 1% BSA, 0.2% gelatin, and 0.05% saponin in PBS and incubated overnight at 4°C with primary antibodies diluted in PBS containing 0.1% BSA and 0.3% Triton X-100. The following day, sections were brought to room temperature, washed, and incubated with fluorescence-conjugated secondary antibodies for 1 h at room temperature in a humidified chamber. Sections were washed, rinsed in distilled water, and mounted with DakoCytomation Fluorescent Mounting Medium.

For quantification of rhodopsin-positive phagosomes, tissue was incubated with primary antibody against rhodopsin (clone 1D4; Abcam, Cat# ab5417; 1:1000), followed by donkey anti-mouse Alexa Fluor 647 secondary antibody (1:600) and DAPI (1:1000).

For quantification of cathepsin D-positive particles, tissue was incubated with primary antibodies against cathepsin D (clone E179; Cell Signaling Technology, Cat# 69854; 1:100), rhodopsin (clone 4D2; Sigma-Aldrich, Cat# MABN15; 1:500), and megalin (abcam cat#ab313741; 1:1000), followed by Alexa Fluor 568-conjugated anti-rabbit (Cat# A-11011; 1:500), Alexa Fluor 488-conjugated anti-mouse (Cat# A-11001; 1:500), and Alexa Fluor 647-conjugated anti-sheep secondary antibodies (Cat# A-21448; 1:500).

### Optical Coherence Tomography

Longitudinal OCT imaging was performed over an 18-month period using the MICRON® IV image-guided OCT 2 system (Phoenix Research Laboratories, Pleasanton, CA, USA), following a protocol adapted from a previous study using the same platform[40]. Horizontal B scans were acquired with the optic nerve head centered in the imaging field. Scans were selected based on image quality and anatomical consistency, and all imaging, segmentation, and analyses were performed blinded to genotype. OCT images were loaded into InSight software (Phoenix Research Laboratories), and retinal layers were manually segmented. Segmented measurements included total retinal thickness, ganglion cell layer/inner plexiform layer, inner nuclear layer/outer plexiform layer, outer nuclear layer and outer segments/RPE. Segmentation coordinates were exported as .csv files and analyzed in R. Measurements were centered relative to the optic nerve head, and values between −100 µm and +100 µm from the optic nerve head center were excluded from analysis. For each retinal layer, the area under the curve (AUC) was calculated across the remaining retinal span to quantify overall structural changes. To reduce inter-animal anatomical variability, AUC values were normalized to baseline measurements obtained three weeks prior to tamoxifen treatment. Longitudinal changes in baseline-normalized AUC values were analyzed using linear mixed-effects models with genotype, time, and genotype-by-time interaction as fixed effects and animal identity as a random effect. Time was treated as a categorical variable. Post hoc comparisons between genotypes at individual time points were performed using estimated marginal means (emmeans) with Holm-adjusted P values for multiple testing correction. The number of animals and eyes included at each time point varied due to animal loss during the study period and transient ketamine-induced cataracts that occasionally prevented reliable OCT imaging.

### RPE Flatmounts

Immediately after euthanasia, eyes were enucleated and fixed in 4% paraformaldehyde (PFA) for 1 hour at room temperature, followed by storage in phosphate-buffered saline (PBS) overnight at 4°C. The following day, eyes were dissected under a stereomicroscope. Connective tissue and the optic nerve were carefully removed using microsurgical scissors. The anterior segment, including the cornea and iris, was excised, and the lens and neural retina were gently removed to expose the underlying RPE layer. Four radial incisions were made from the peripheral edge toward the optic nerve head to create four quadrants, allowing the eyecup to be flattened for immunohistochemistry preparation. The RPE flatmounts were stored in 1% PFA until immunostaining.

### RPE flatmount immunofluorescence microscopy

For F-actin cytoskeleton and RPE protein labeling, RPE flatmounts were washed in 2X PBS and blocked in blocking buffer (2.5% BSA, 3% Triton-X-100, 0.5% Tween in PBS) for 1 hour at room temperature on a rocking table. RPE flatmounts were immunolabeled overnight at room temperature with primary antibodies recognizing megalin (1:1000). Following washing in blocking buffer, AlexaFluor-conjugated secondary antibodies, F-actin cytoskeleton stain (Life Technoliges, CA Cat#R415), and DAPI nuclei stain were applied for 2 hours at room temperature. The tissue was washed in 2X PBS and mounted with Dako Fluorescence Mounting Medium.

Fluorescence images were acquired using a slide scanner (Olympus VS120) for quantification of knockout efficiency and a laser-scanning confocal microscope (710 or 800 laser scanning confocal microscope, Zeiss) for visualization of cell morphology. The confocal images were captured using 63x/1.4 oil Plan-Apochromat and subsequently processed in ImageJ/FIJI. For slide scanner imaging, 40x magnification and Maximum Intensity Projection (Z-range 10 µm, Z-spacing 2 µm, 5 planes) were employed. For improved visualization of F-actin-defined cell borders, the Olympus deblur function was applied. The images were processed and analyzed using QuPath software and cell segmentation was performed using the BIOP qupath-extension-cellpose plugin[41]. Cells were classified as either megalin^+^ cells or megalin^-^ cells by manual thresholding, and the cell area (µm^2^) was measured in QuPath and further analyzed in R Studio.

### Cell size analysis

RPE flatmount images were obtained from 8 animals (4 WT, 4 KO). Individual RPE cells were segmented and morphological features extracted, including cell area, length, circularity, solidity, and diameter measurements using CellPose. Because thousands of cells were measured per animal, the data had a hierarchical structure with cells nested within animals and animals nested within experimental groups. Cells with missing or undefined classification labels were excluded prior to analysis. To avoid pseudoreplication, statistical analyses accounted for the non-independence of cells from the same animal.

For descriptive purposes, summary statistics were calculated both at the group level (all cells pooled) and at the animal level, including mean cell area, standard deviation (SD) of cell area, and number of analyzed cells per animal. Cell area distributions were assessed visually using Q–Q plots and showed deviation from normality. Differences in mean cell area between WT and KO were therefore analyzed using a linear mixed-effects model with group as a fixed effect and animal identity as a random effect (Area ∼ Group + (1 | Image)), fitted by restricted maximum likelihood (REML).

To assess differences in variability, a mixed-effects model allowing group-specific residual variance was fitted using a varIdent variance structure (∼1 | Group) and compared with a model assuming equal variance using a likelihood ratio test. As a complementary conservative analysis, per-animal mean and SD of cell area were compared between groups using Wilcoxon rank-sum tests, treating each animal as an independent biological replicate. Data were visualized using violin plots for cell-level distributions and beeswarm plots for animal-level summaries, with group mean ± SD overlaid. All analyses were performed in R using lme4, nlme, ggplot2, and ggbeeswarm.

### Rhodopsin analysis

Paraffin-embedded eyes from WT and KO mice one month after induction were staining with rhodopsin antibody (Sigma-Aldrich cat#MABN15) and imaged using a slide scanner (Olympus VS120) with 40x magnification and Maximum Intensity Projection (Z-range 10µm, Z-spacing 2µm, 5 planes). The posterior half of the RPE was marked manually in QuPath and the images were exported to ImageJ for particle analysis. For all images, brightness was adjusted to 80-4000 and the threshold was set to 53 for quantification. Counts were normalized the RPE length to account for differences in the size of the ROI.

### Quantification of cathepsin D-positive particles

Paraffin-embedded retinal cross-sections containing RPE/choroid only were immuno-labeled for cathepsin D, rhodopsin, and megalin. Paraffin sections were imaged on a DeltaVision OMX SR microscope (GE Healthcare) equipped with an Olym-pus PlanApo N 60×/1.42 NA oil immersion objective and PCO.edge sCMOS cameras. Images were acquired in conventional widefield mode using sequential excitation at 488 nm (20 ms exposure), 568 nm (100 ms), and 640 nm (150 ms), with emission collected at 527, 603, and 679 nm, respectively. Immersion oil with a refrac-tive index of 1.522 was used. Z-stacks were collected at a step size of 0.125 μm, with the number of optical sec-tions adjusted to span the full sample thickness, and a lateral pixel size of 0.08 μm (512 × 512 pixels per plane). Raw image stacks were deconvolved and channel-aligned in softWoRx 7.2.2 (GE Healthcare).

Multichannel z-stacks were batch-processed in Fiji/ImageJ using a custom macro. Maximum intensity projections of the cathepsin D channel were generated, and particles were identified by fixed intensity thresholding followed by binary mask segmentation. Particles touching image borders were excluded, and particle count and area were quantified using the Analyze Particles function. Quality control masks were generated for all analyzed images. Particle counts were normalized to manually measured RPE length for each image and expressed as particles per µm RPE.

Statistical analyses were performed in R Studio using the lme4, lmerTest, effsize, and ggplot2 packages. Because multiple images were obtained from each animal, differences between WT and KO animals were analyzed using a linear mixed-effects model with group as a fixed effect and animal identity as a random effect (ParticlesPerUm ∼ Group + (1 | AnimalID)). Animal-level means were additionally com-pared using Welch’s t-test. Normality was assessed using the Shapiro–Wilk test, and effect size was estimated using Cohen’s d. For figure preparation only, background subtraction was performed in Fiji using a rolling ball radius of 50 pixels, and fluorescence intensities were adjusted identically across WT and KO images within each channel.

### Cathepsin D activity assay

RPE cells were isolated using the same protocol as for Western blotting. Cells were lyzed and Cathepsin D activity was measured using a fluorometric activity kit from abcam cat#65302 (Abcam, UK) according to the manufacturer’s instructions.

### Statistical analysis

Unless otherwise stated, statistical analyses were performed using unpaired two-tailed Student’s t-tests for two-group comparisons and one-way ANOVA with post hoc multiple-comparison correction for analyses across time points.

## Acknowledgements & Disclosures

We thank Pia K. Nielsen, Helle Salling Gittins, Hanne Sidelmann and Cassandra Bennetzen from Aarhus University for technical assistance. We thank Dr. Steven Tsang from Columbia University for supply of bovine photoreceptor outer segments, and Dr. Pierre Verroust for supply of the sheep-anti-megalin antibody. We also thank David S. Williams (Stein Eye Institute, UCLA) for providing antibodies and equipment and for access to the microscope used to acquire the images in Fig. 4A, and Antonio E. Paniagua for assistance with the microscopy.

We thank the Laboratory Animal Core Facility, the Bioimaging Core for imaging assistance, and the Phenotyping Core Facility, Department of Biomedicine, Health, Aarhus University, Denmark, for the use of equipment and support. Imaging was supported by the Novo Nordisk Foundation Large Infrastructure grant NNF24OC0088843. The mass spectrometry equipment utilized in this study is part of the Danish Single-Cell Examination Platform (CellX) established with support from the Danish Research Agency Infrastructure Program (5229-0009B).

## Author Contributions

VBM and RN had full access to all the data in the study and takes responsibility for the integrity of the data and the accuracy of the data analysis. Conceptualization: DKR, PLM, SHL, VBM, RN. Data curation: DKR, PLM, SHL, TS, QW. Formal analysis: DKR, PLM, SHL, VBM, RN. Visualization: DKR, PLM. Resources: SHL, TSJ, ALA, QW, RAF, VBM, RN. Writing – original draft: DKR, PLM, VBM, RN. Writing – review and editing: All authors. Funding acquisition: VBM, RN. Supervision: VBM, RN.

## Institutional Review Board Statement

The study was approved by the Institutional Review Boards at Stanford University and adhered to the tenets set forth in the Declaration of Helsinki.

## Informed Consent Statement

Not applicable.

## Data Availability Statement

The mass spectrometry proteomics data have been deposited to the ProteomeXchange Consortium via the PRIDE partner repository with the dataset identifier PXD079688. All other original data is either available in the supplementary material or will be provided by the lead contact upon request.

## Declaration of generative AI and AI-assisted technologies

During the preparation of this work, the authors used ChatGPT to provide feedback on writing and coding. After using this tool/service, the authors reviewed and edited the content as needed and take full responsibility for the content of the published article.

