## Supplemental materials for "Megalin deficiency perturbs retinal homeostasis and impairs cathepsin D processing and phagosome-lysosome maturation in the retinal pigment epithelium"

**Supplementary Online Content**

**Table of Contents**

**Supplemental Figures**

**Figure S1.** RPE morphology

**Figure S2.** Retinal histology

**Figure S3.** Axial length measurements

**Figure S4:** Proteomic analysis

**Figure S5.** Full Western blots

**Supplemental tables**

**Table S1.** Aptamer-based proteomics human iPSC-RPE

**Figure S1. RPE morphology**

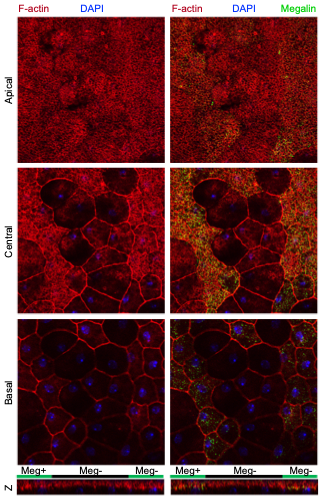

RPE morphology at 3 month after induction. Apical, central and basal stainings reveal altered RPE morphology in megalin-KO mice.

**Figure S2: Retinal histology**

**A)**

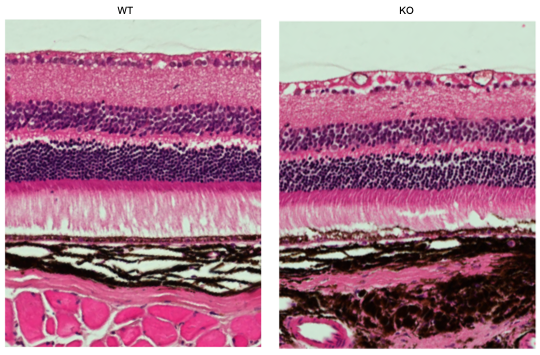

**B)**

A) Retinal histology of a representative WT mouse and the KO mouse with the lowest degree of retinal degeneration after 12 months are shown. B) time course of HE-stainings showing limited change over the first 6 months.

**Figure S3: Axial length measurement**

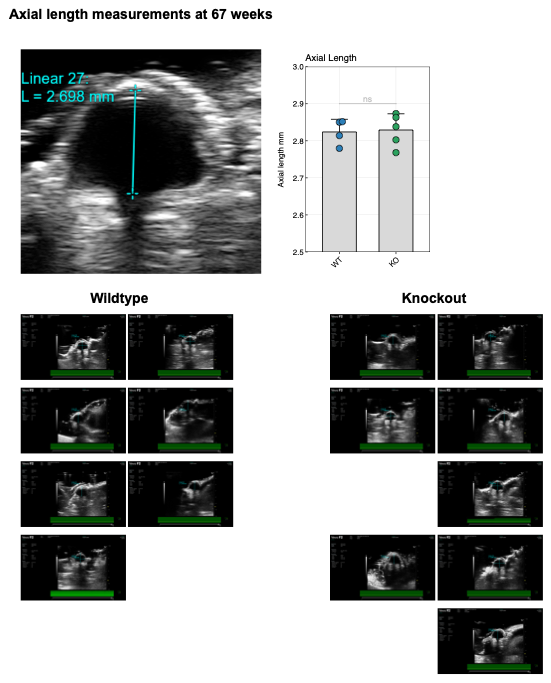

**Figure S4: Proteomic Analysis**

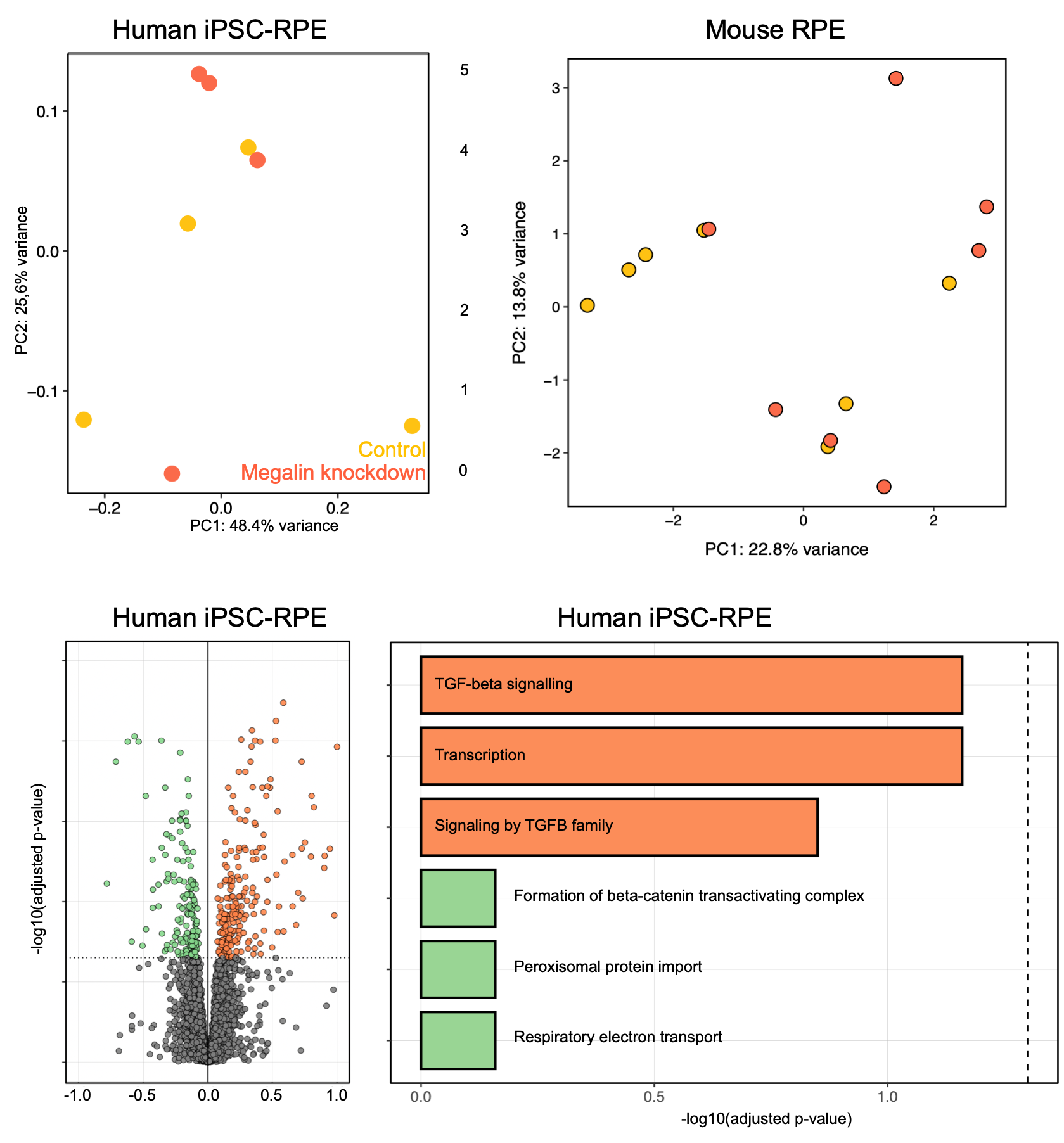

Principal component analysis of human iPSC-RPE and mouse RPE cells comparing KO with WT reveals partial separation. Volcano plot and unsupervised pathway analysis of human iPSC-RPE reveals no significantly altered pathways, but most prominent signal in TGF-beta signaling and transcription pathways.

**Figure S5: Original Western Blots**

*Characterization of iPSC-RPE*

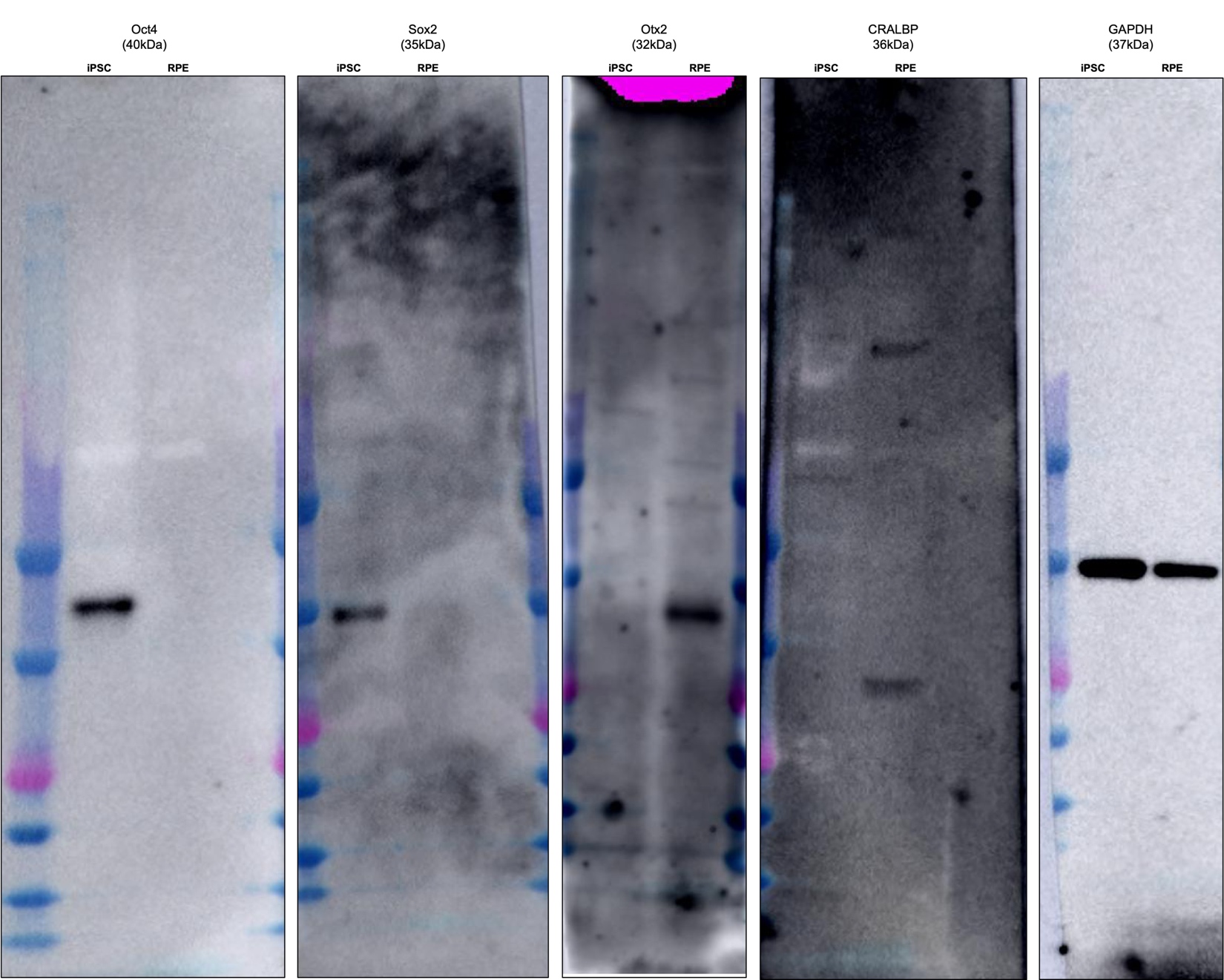

*In Vitro Rhodopsin accumulation*

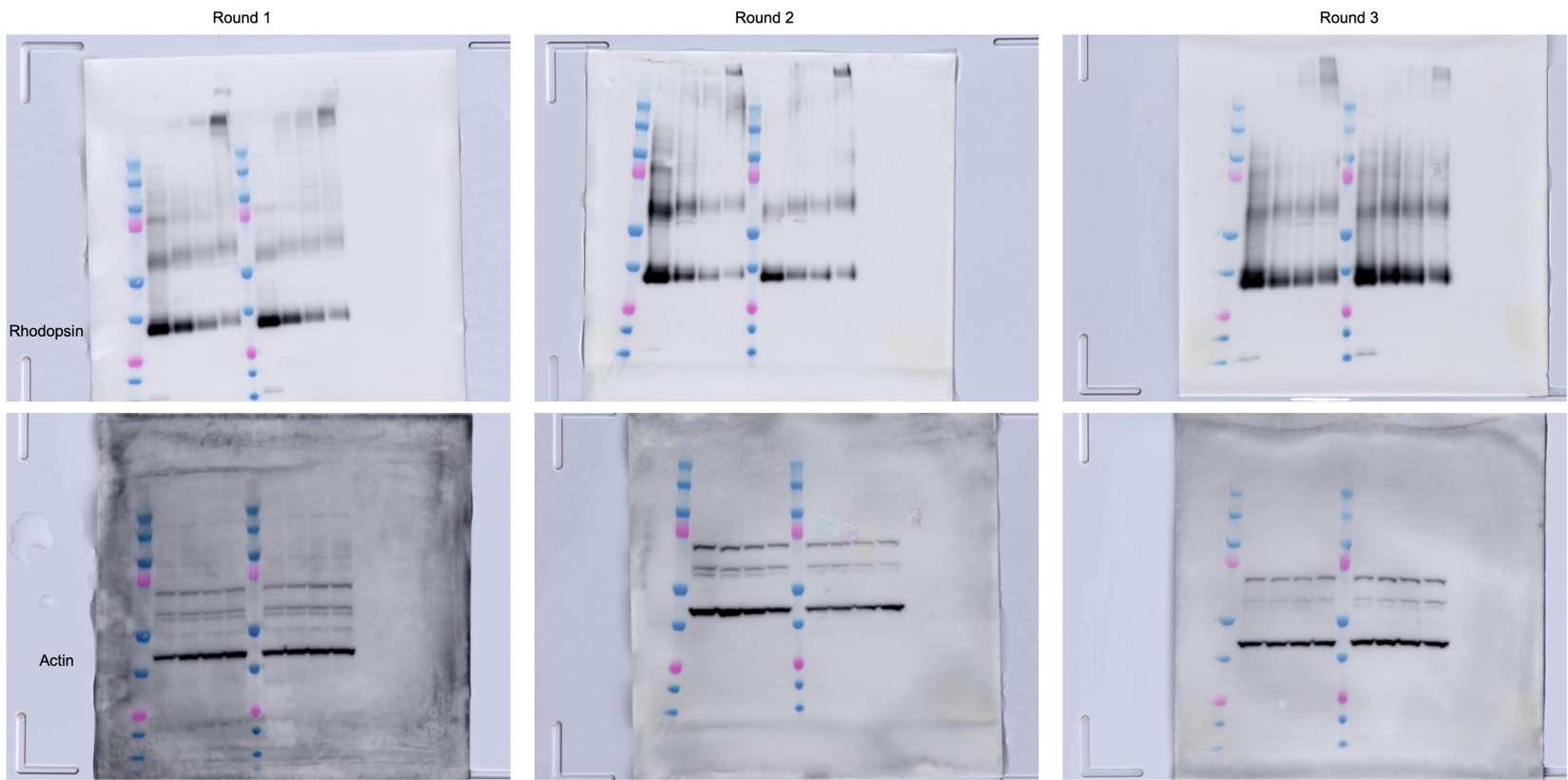

*In Vivo Cathepsin D*

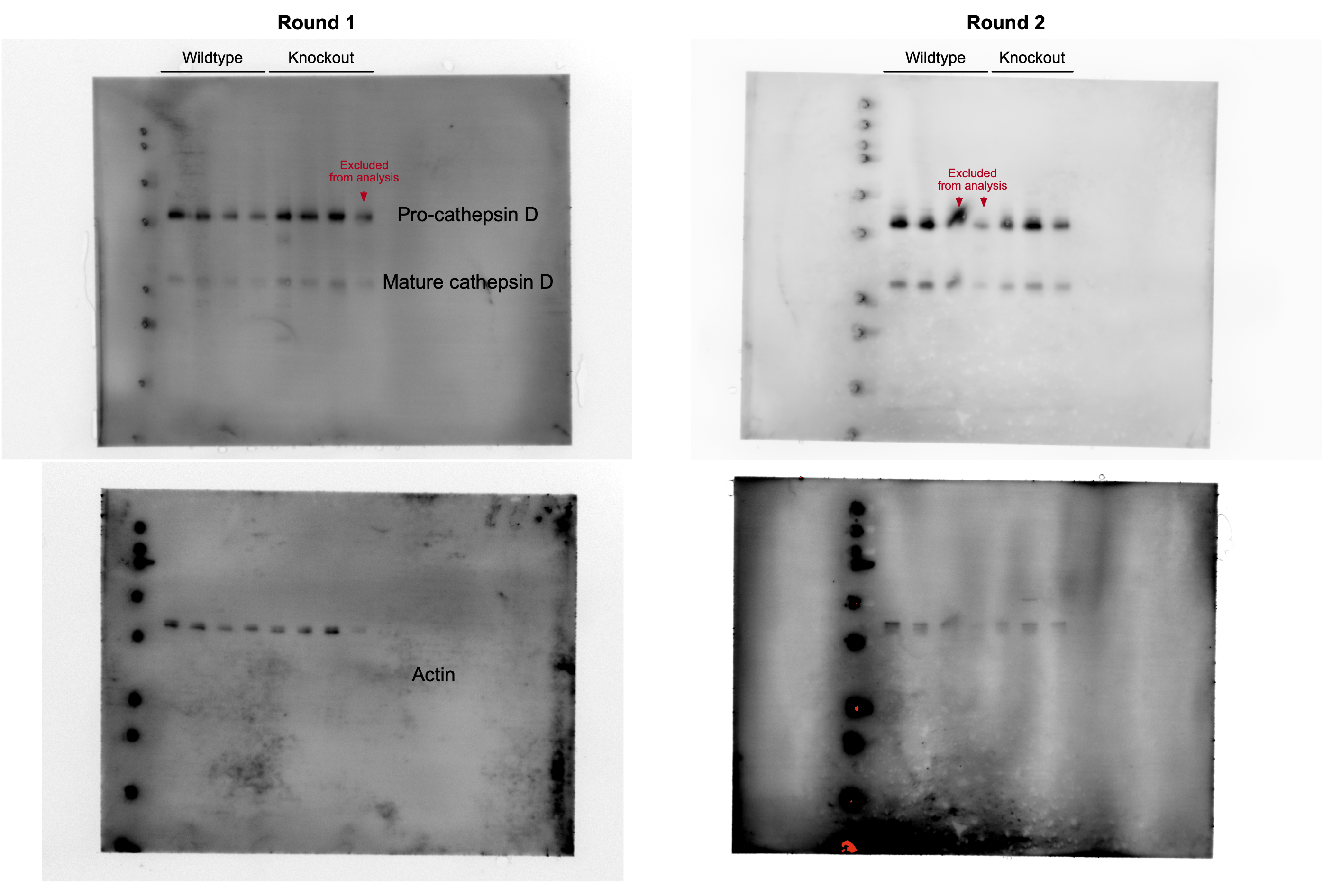
